# Comparing Imaging Biomarkers of Cerebral Edema after TBI in Young Adult Male and Female Rats

**DOI:** 10.1101/2021.06.25.449932

**Authors:** Heather M Minchew, Sadie L Ferren, Sarah K Christian, Jinxiang Hu, Paul Keselman, William M Brooks, Brian T Andrews, Janna L Harris

## Abstract

Traumatic brain injury (TBI) is one of the leading causes of death and disability worldwide. Cerebral edema following TBI is known to play a critical role in injury severity and prognosis. In the current study we used multimodal magnetic resonance imaging (MRI) to assess cerebral edema 24 hours after unilateral contusive TBI in male and female rats. We then directly quantified brain water content in the same subjects *ex vivo*. We found that both males and females had similarly elevated T_2_ values after TBI compared with sham controls. Apparent diffusion coefficient (ADC) was more variable than T_2_ and did not show significant injury effects in males or females. Brain water was elevated in male TBI rats compared with sham controls, but there was no difference between female TBI and sham groups. Notably, MRI biomarkers of edema were more closely correlated with brain water in male rats; female rats did not show any relationship between brain water and T_2_ or ADC. These observations raise questions about the interpretation of radiological findings traditionally interpreted as edema in female TBI patients. A better understanding of sex differences and similarities in the pathophysiology of post-traumatic edema is needed to help improve patient management and the development of effective treatment strategies for men and women.

## INTRODUCTION

Traumatic brain injury (TBI) is a leading cause of death and disability in the United States^1,2^. The initial physical impact of TBI leads to the initiation of a complex molecular and cellular cascade of secondary injury mechanisms, among which cerebral edema is a critical contributor. Post-traumatic edema is strongly predictive of poor patient outcomes following TBI^3-6^.

Cerebral edema, defined as an elevation of brain tissue water, has traditionally been categorized as either vasogenic or cytotoxic in origin. More recently it has been proposed that this distinction may be somewhat arbitrary since the molecular mechanisms of both processes are highly inter-related^7^. Vasogenic edema arises from direct disruption of the blood brain barrier and leaking of solute-laden fluid from the blood into the extracellular space of the brain. Cytotoxic edema results from disruptions to cell membrane ion channels, causing ion-driven movement of water into cells. Quantitative diffusion-weighted imaging (DWI) and T_2_ mapping have each been used as magnetic resonance imaging (MRI) biomarkers of cerebral edema^8-12^. The apparent diffusion coefficient (ADC) calculated from DWI is a measure of water diffusion within the target tissue, which is influenced by the relative balance of water in the intracellular versus the extracellular compartment. Thus, elevated intracellular water (i.e. cytotoxic edema) tends to lower the ADC values, while elevated extracellular water (vasogenic edema) tends to increase ADC^12-16^. By contrast, T_2_ mapping measures the transverse relaxation time of protons and is sensitive to the total water content in the tissue regardless of cellular compartment.

Although women incur a significant number of TBIs, much is still unknown regarding differences in pathophysiology between men and women^17,18^. Historically, men have outnumbered women in clinical trials and most studies in animal models have been limited to male animals^17,18^. The severity, type, and duration of symptoms appear to differ between men and women who have experienced a TBI, although studies to date have not supported a significant association between sex and injury-induced mortality ^19-23^. In rodent TBI models, sex differences in brain damage and neurobehavioral recovery have long been established^24^. Steroid hormones such as estrogen and progesterone are thought to play a protective role against brain edema development, a factor which may help explain significant sex differences^25-27^. Clinically, cerebral edema presents as an increase in intracranial pressure (ICP), a measure significantly associated with increased morbidity and mortality^3,6,28^. Sex hormones may influence ICP; in particular the high circulating estrogen and progesterone levels in women during the proesterous stage of the reproductive cycle may attenuate edema and ICP, conferring neuroprotection^26,29^. However, it was also reported that female rats that were ovariectomized (non-cycling) had less edema after TBI than both cycling females and male rats, suggesting that non-hormonal factors may also contribute to sex differences in cerebral edema^30^.

Previous studies in animal models have used the “wet-dry” method to characterize cerebral edema in diffuse and focal TBI models^31-34^. This *ex vivo* approach directly quantifies the tissue water content in the brain, where freshly dissected tissue is weighed on a fine scale, desiccated, and weighed again to calculate the percent tissue water. Translation of findings to humans, however, requires alternative non-invasive methods. MRI is routinely used for clinical management in stable TBI patients^35-37^. MRI is also increasingly being used in animal models to investigate TBI mechanisms including cerebral edema. However, to date no studies have used MRI methods to specifically interrogate possible sex differences in edema after TBI. Therefore, the goal of this study was to compare cerebral edema in male and female rats after moderate contusive TBI using multimodal MRI followed by direct *ex vivo* quantification of brain water content.

## RESULTS

### T_2_ -weighted image mapping

T_2_ measurements in an ROI centered on the injured cortex and a homologous control ROI on the contralesional side (**Fig. 1A**) were used to calculate the change in T_2_ for each animal. Two-way ANOVA showed a main effect of group (F(1,36) = 29.38, p < 0.0001) indicating a significant effect of TBI. There was no main effect of sex and no interaction, indicating no significant difference in T_2_ between males and females. Testing within each sex showed the change in T_2_ was significantly higher in the male TBI group compared with male sham controls (p < 0.0001), and in the female TBI group compared with female sham controls (p = 0.001) (**Fig. 1B**).

**Figure 1.**
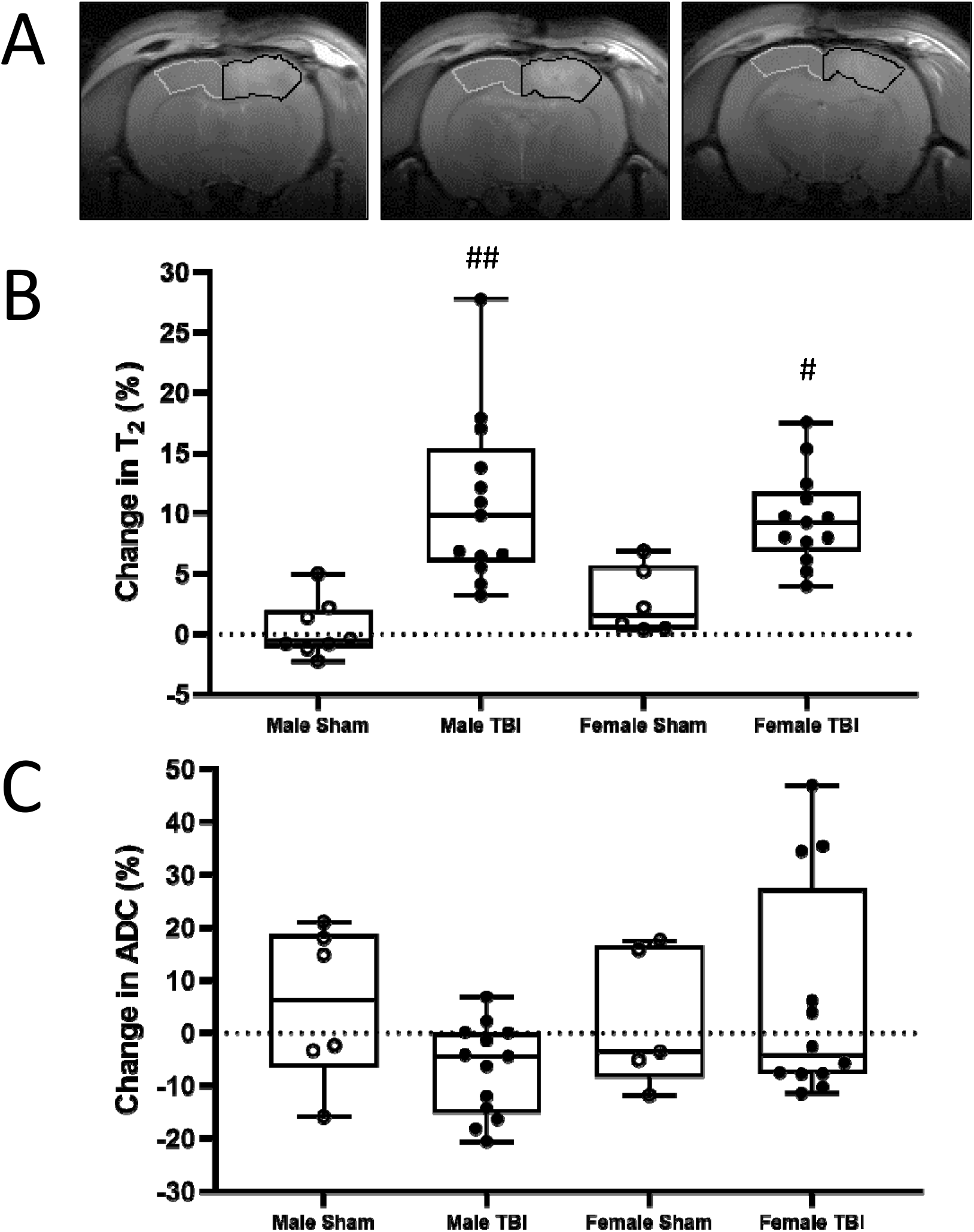
Magnetic resonance imaging measures of cerebral edema after TBI in male and female rats. **(A)** Regions of interest (ROI) for MRI analysis are shown on three sequential diffusion weighted images encompassing the center of the injury in a male rat 24 hours after controlled cortical impact. Contralesional ROI = grey outline, injured ROI = black outline. The T_2_ and ADC values were averaged across the three slices. **(B)** Plot shows the calculated change in T_2_ in the injured ROI compared with the contralesional side for each subject. **(C)** Plot shows the calculated change in apparent diffusion coefficient (ADC) in the injured ROI compared with the contralesional side for each subject. # p < 0.05, ## p < 0.001 compared to sex-matched sham group.

### Diffusion weighted imaging

ADC measurements from the same injured and contralesional ROIs were used to calculate the change in ADC for each animal. Change in ADC was more variable than T_2_ at 24h after TBI, with some individuals increasing on the injured side and others decreasing in all groups. Two-way ANOVA showed no main effect for group or sex, and no significant two-way interaction TBI (**Fig. 1C**). Although there were no main effects, since previous studies generally restricted to males have found reduced ADC acutely after TBI^11,14,38,39^, we separately explored effects of injury on ADC within each sex. The change in ADC was not significantly different in the male TBI group compared with male sham controls (p = 0.11) or in the female TBI group compared with female sham controls (p = 1.0).

### Brain water content

Water content of the injured brain tissue sample and the contralesional sample were used to calculate the change in brain water content. Brain water changes were somewhat variable in all groups; two-way ANOVA showed no main effect of group or sex, and no significant two-way interaction. Since previous studies, most often restricted a single sex, have found elevated brain water acutely after TBI^30,33,34,38,40^, we separately compared brain water in the TBI and sham groups within each sex. In males, tissue water was elevated in the TBI animals compared with the sham controls (p = 0.033). In females, the change in brain water was not different between the TBI group and the sham controls (p = 0.83) (**Fig. 2**).

**Figure 2.**
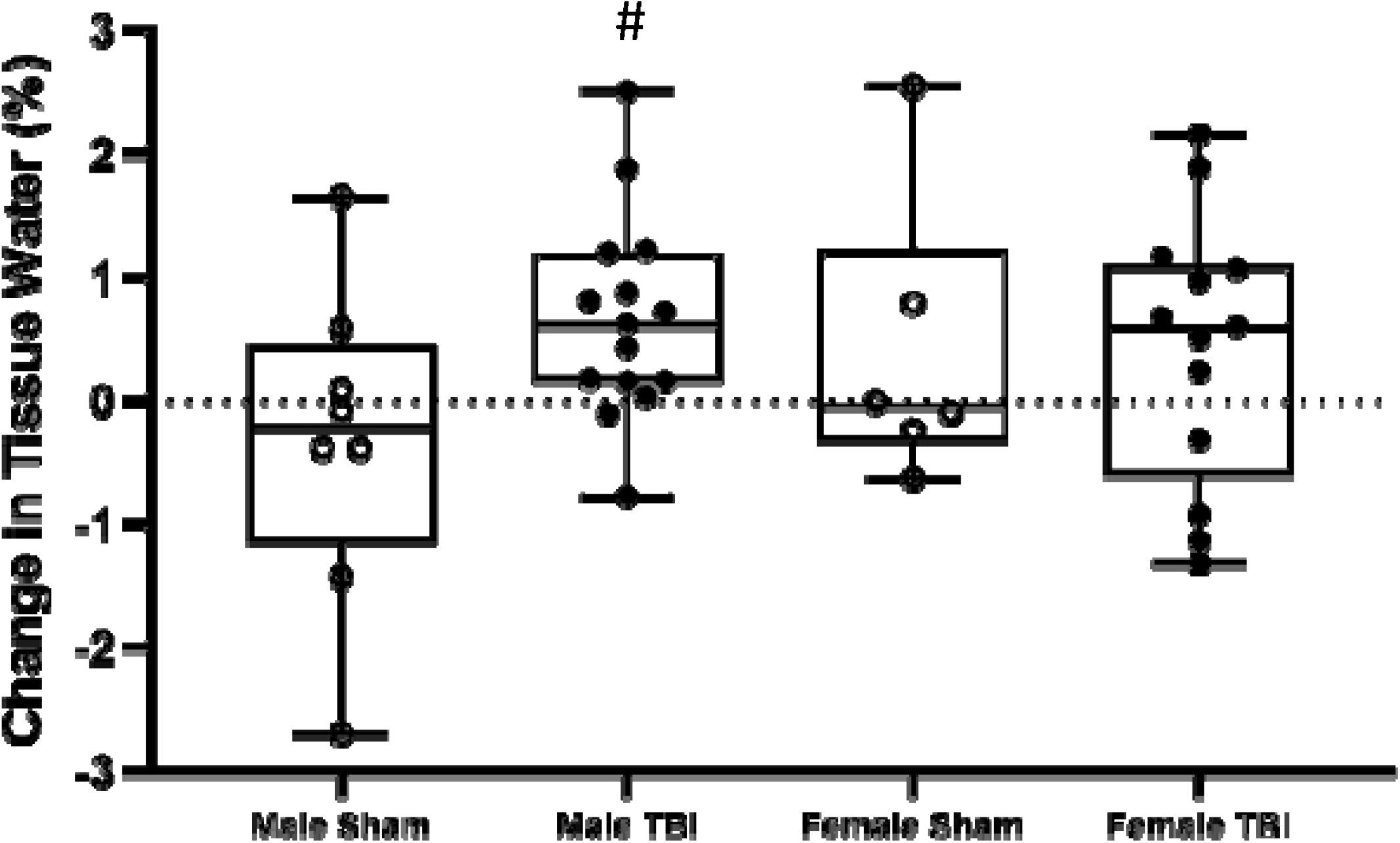
Change in brain water after TBI in male and female rats. Plot shows the calculated change in brain water content in the injured tissue sample compared with the contralesional side for each subject. # p < 0.05 compared to sex-matched sham group.

### Relationship between brain water content and imaging measures of edema

In the injured hemisphere of male rats, there was a positive correlation between brain water values and T_2_ values (r = 0.65, *p* = 0.02; **Fig. 3**). In addition, brain water in male rats trended toward a negative correlation with ADC values, but this did not reach significance (r = - 0.44, *p* = 0.13). In female rats the brain water content in the injured hemisphere was not significantly correlated with either T_2_ values (r = 0.20, *p* = 0.53) or with ADC values (r = -0.22, *p* = 0.52).

**Figure 3.**
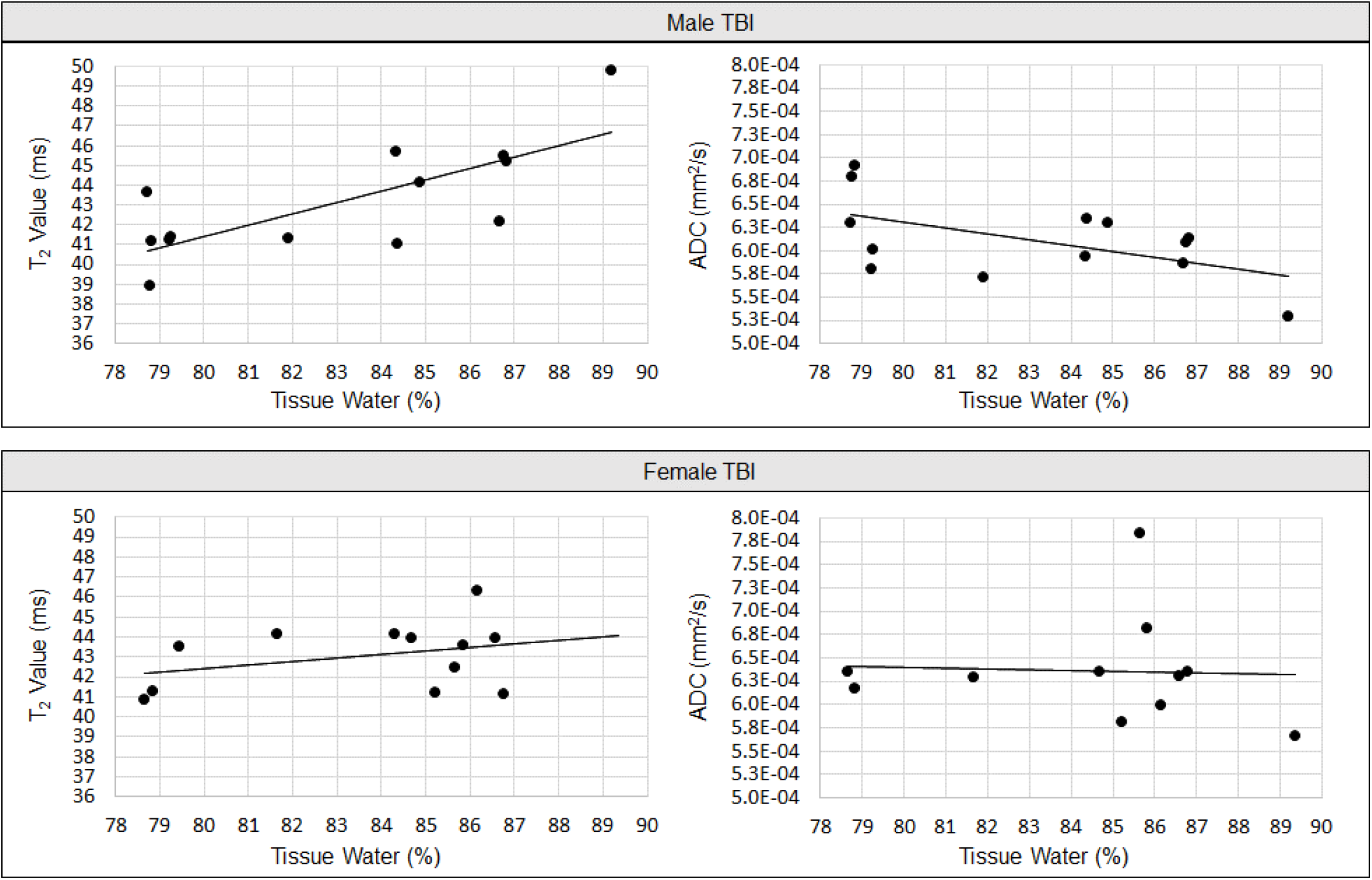
Relationship between brain water content and MRI measures of cerebral edema after TBI in male and female rats. In the injured hemisphere of male rats, tissue water was positively correlated with T_2_ values (r = 0.65, *p* < 0.05) and showed a trend toward negative correlation with ADC that did not reach significance (r = -0.44, *p* = 0.15). By contrast, in the injured hemisphere of female rats, water content was not significantly correlated with T_2_ values (r = 0.20) or with ADC (r = -0.22). Note that some animals with partial data were included in the study (e.g. successful T_2_ but not DWI scan, see details in Methods).

## DISCUSSION

Sex differences in TBI damage and subsequent functional recovery have long been observed. However, the mechanisms underlying this observation have not been well characterized. This study used MRI biomarkers and brain water measurements to assess cerebral edema 24 hours after contusive TBI in male and female rats. At this acute time point, cortical T_2_ values were similarly elevated in the male and female TBI groups, but not in the sham controls. Change in ADC was more variable, with some individuals showing increased ADC and others decreased ADC after the TBI/sham procedure in all groups. In males and females, we found no significant difference between TBI and sham groups for ADC. Taken together, one interpretation is that T_2_ mapping is a more sensitive measure of cerebral edema than diffusion weighted imaging. Alternatively, ADC could reflect a more heterogeneous manifestation of edema, consistent with the variability previously reported for ADC measured after TBI in male rats^9^. This variability could be attributed to the concurrent mix of vasogenic/cytotoxic edema developing over a time course that may vary between individuals, even in a controlled animal study.

Translational neuroimaging studies in animals allow for direct comparison between non-invasive MRI biomarkers and invasive assays to confirm pathophysiological mechanisms. We found that change in brain water content was somewhat variable, and when brain water was compared across all groups there was no significant effect of injury or sex. Since previous rat TBI studies, typically limited to a single sex, have shown elevated brain water at 24 hours, we separately explored possible injury effects within each sex. This approach revealed increased brain water in male TBI rats compared to sham controls, but no difference between female TBI and sham. Additionally, we explored the relationship between *in vivo* MRI and *ex vivo* tissue water measured in the same subjects. We found that T_2_ and ADC values—imaging measures that are used to make clinical inferences about cerebral edema—more closely correlated with brain water measures in male rats than in females after TBI. The expected relationships between MRI and confirmed mechanism were observed in the males: higher brain water correlated with higher T_2_ values, and trended toward correlation with lower ADC values (although this did not reach significance). By contrast, female rats did not show any clear relationships between brain water and MRI biomarkers. The reason for this is not currently clear, but the observation raises important questions about clinical interpretation of radiological findings traditionally interpreted as biomarkers of edema, when observed in female TBI patients. Future larger studies are needed to confirm this possible sex difference in imaging biomarkers of edema, and to explore mechanisms that may modulate the relationship between elevated brain water and MRI measures.

In this initial exploratory study we did not track cycle stage or control for hormonal status in female rats, since in the real world women may sustain TBI at any time in their cycle. A growing body of research indicates that estrogen and progesterone may attenuate certain aspects of TBI pathophysiology, including edema^26,27,41-45^. It is possible that differing sex hormone levels associated with cycle stage contributed to the greater variability in ADC that we observed in female rats compared with males. Future studies are warranted to investigate this possibility. However, we also observed approximately equal variability of T_2_ and brain water measurements in males and females. This underscores the growing recognition that hormonal effects on TBI outcomes are complex and may depend on the particular outcome measure used, the time point, and characteristics of the injury^17,18,26,46-49^.

We measured cerebral edema 24 hours post injury, a time point that corresponds to maximal edema after CCI in adult male rats^33,34,38,50^. Since the development of edema after brain trauma is a dynamic process, the timing of which is likely to be affected by the severity and type of injury, results from single time point studies should be interpreted with caution. A possible interpretation of our tissue water results is that females develop less severe edema than males after identical TBI procedures; an alternative explanation might be that the time course of edema development differs by sex. Indeed, in a time course study of diffuse brain trauma, O’Connor et al. reported that female rats had more severe edema than males at 24h, *and also* had less edema than males at earlier and later timepoints^51^. To date, sex differences in the time course of edema have not been investigated after focal TBI, and longitudinal studies that include both sexes remain an important area for future research.

The surgeries and imaging procedures in this study were performed under isoflurane anesthesia, with the dose for each animal adjusted during the course of each procedure based on physiological parameters. If male and female rats received different anesthesia doses this could affect results, so we reviewed our study records to calculate the anesthesia duration and the cumulative anesthesia dose (percent isoflurane x time) for each animal. We found no differences between males and females in either measure (**Supplemental Table 1**). Previous studies have reported that single or repeated isoflurane exposure can lead to various cognitive and pathophysiological changes in the brain ^52,53^. In particular, there is evidence that isoflurane may worsen post-traumatic edema, but these studies were conducted exclusively in male animals^40,54,55^. The effect of isoflurane on cerebral edema in females is unknown, but if it is similar to males then we would expect the similar doses we administered to affect both sexes equally. We also used bupivacaine for local analgesia; because this was administered extracranially at the incision site we believe it unlikely to have affected edema within the brain. Little is known about possible sex differences in bupivacaine effects, but one study reported a higher potency in women than men after intrathecal administration^56^. The requirement for anesthetics and analgesics is an inherent limitation of invasive studies in animal models, but future studies can help address this by intentionally matching the dose between experimental groups, tracking dosage for use as a covariate, and considering alternative analgesic/anesthetic agents depending on study goals^38,50,51,57^.

Due to a historic focus on males in TBI research much remains unknown about sex differences in TBI pathophysiology, and the relative lack of data on the time course and molecular mechanisms of cerebral edema in females needs to be addressed. A greater number of studies specifically comparing males and females will lead to better understanding of the differences and similarities between sexes, ultimately driving improved patient management and the development of effective treatment strategies.

## METHODS AND MATERIALS

### Animals

Twenty-three male and 23 female rats (F344; Charles River, Wilmington, MA, USA) were used in the study. Rats were 67-83 days old and weighed 157-229 g (male) and 112-152 g (female) at the time of surgery. Animals were housed in same-sex pairs on a 12-hour light-dark cycle with free access to food and water. All protocols were approved by the University of Kansas Medical Center Animal Care and Use Committee consistent with the standards of animal care in the United States Public Health Service Policy on the Humane Care and Use of Laboratory Animals.

### Controlled Cortical Impact (CCI) and Sham surgeries

All surgeries were carried out by a single surgeon, following methods previously reported^58-60^. Each surgery cohort of 4-8 rats contained an equal number of males and females. Surgeries were carried out in randomized order; surfaces were cleaned between procedures to minimize potential stress from exposure to the opposite sex. For surgery, animals were anesthetized with isoflurane (4% induction, 2% maintenance in 2:1 medical air:oxygen) and body temperature was maintained with a heating pad. The head was immobilized in a stereotaxic frame, shaved, and scrubbed with iodine and ethanol. Bupivacaine (0.25%) was administered subcutaneously at the surgical site. Utilizing aseptic technique, a midline incision was made and the skull exposed. The circular craniectomy was formed by a 6mm Michele trephine over the right sensorimotor cortex, lateral to bregma and centered between bregma and the temporal ridge. Moderate CCI was delivered with an impactor mounted to a stereotaxic arm (Leica; Saint Louis, MO; impactor tip diameter = 5mm; velocity = 5 m/s; depth = 2mm; contact time = 300 ms; angle = 5° from vertical). Sham rats received the same procedure without impact; anesthesia time for sham surgeries was extended to match the average time for CCI surgeries. The incision was sutured closed and animals were transferred to a heated recovery cage designated for a single sex. After recovery of locomotion, animals were returned to their home cages.

### Magnetic Resonance Imaging (MRI)

MRI scans were performed approximately 24 hours after TBI (average 23.4 ± 2.8 hours) on a 9.4 Tesla system with a Varian INOVA console (Agilent Technologies, Santa Clara, CA). The system is equipped with a 12 cm gradient coil (40 G/cm, 250 μs; Magnex Scientific, Abingdon, UK). A custom-made quadrature surface radiofrequency coil, consisting of two geometrically decoupled 18mm loops, was placed on the animal’s head to transmit and receive at 400 MHz.

During imaging, anesthesia was delivered via nosecone (1.5 - 3% isoflurane in 2:1 medical air:oxygen) to maintain a respiration rate of 40 - 80 cycles/minute. Respiration was monitored with a pressure pad (SA Instruments, Stony Brook, NY, USA). Animals were placed on a heating pad and body temperature was maintained at 37 ± 1 °C via feedback control (Cole Parmer, Veron Hills, IL, USA). Transverse and sagittal GEMS localization images were acquired to check the animal’s positioning in the magnet (FOV_ax_ = 2.56 × 2.56 cm^2^, FOV_sag_ = 5.12 × 2.56 cm^2^, matrix = 128 × 128, TR/TE = 90 ms/2.8 ms, slices = 9, thickness = 1 mm). High-resolution T_2_-weighted images were acquired using a RARE sequence (FOV = 2.56 × 2.56 cm^2^, matrix = 256 × 256, TR/TE = 4000/72 ms, averages = 2, thickness = 1 mm, echo train length = 8, echo spacing = 18 ms, total acquisition time ∼4 minutes).

For mapping T_2_ weighted images, a series of multi-slice spin echo acquisitions were performed (FOV = 2.56 × 2.56 cm^2^, matrix = 128 × 128, TR = 3 s, TE = 20, 30, 50, 75, 100, 150 ms, averages = 2, slices = 9, thickness = 0.8 mm with a 0.2 mm gap, acquisition time = 6 minutes 25 seconds for each TE). T_2_ values were calculated for each voxel in MATLAB from a 2-parameter exponential fit of the TE values. The DWI and T_2_ maps were acquired with identical slice positions, FOV, and spatial resolution.

DWI was performed using a standard mono-polar diffusion weighted spin echo sequence (FOV = 2.56 × 2.56 cm^2^, matrix = 128 × 128, TR/TE = 1500/26 ms, gradients applied along readout direction, b = 50, 500, 1000 s/mm^2^, averages = 1, slices = 9, thickness = 0.8 mm with a 0.2 mm gap, total acquisition time 9 minutes 40 seconds). Apparent diffusion coefficient (ADC) values were calculated for each voxel in MATLAB (Mathworks Inc., Natick, MA, USA) from a 2-parameter exponential fit of the b values.

### Brain water content

Immediately following MRI, rats received an overdose of isoflurane and were euthanized via decapitation. Brain tissue was rapidly extracted from the skull, placed in a coronal slicing matrix, and a 10 mm block was cut beginning ∼1 mm rostral to the cortical injury extending caudally (**Fig. 4A**). The block was removed, cut in half horizontally to separate dorsal from ventral structures, then cut in half vertically at the midline (**Fig. 4B**). The dorsal brain samples from the left (contralesional) and right (injured) hemispheres were used for wet-dry analyses. Each sample was immediately weighed (Mettler Toledo AE100 Scale, Columbus, OH, USA) and values were recorded as the wet weight. The samples were then placed in a laboratory oven (Grieve Laboratory Oven Model L R-271C, Round Lake, IL, USA) at 100 °C. Dry weights were recorded after 24, 48 and 72 hours and the lowest dry weight was used in water content calculation; for most subjects this was at 72 hours. Tissue water content was calculated from the wet and dry weights: [wet weight (g) - dry weight (g)] / [wet weight (g)] x 100. Finally, for each animal we calculated the change in tissue water observed on the injured vs the contralesional side: [(Injured side/Contralesional side)*100 – 100].

**Figure 4.**
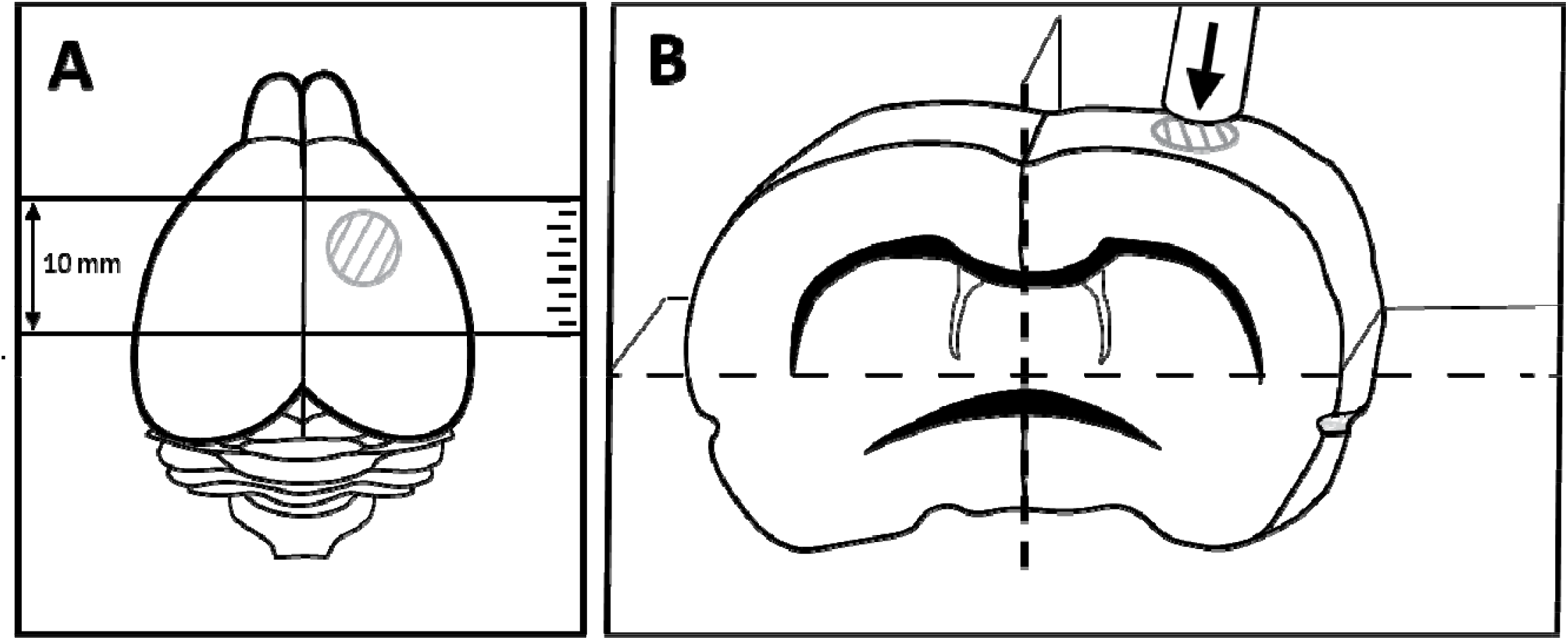
Brain injury location and regions dissected for brain water content analysis. A) dorsal view of the rat brain. The circle depicts the site of craniectomy and cortical impact. Using a coronal slicing matrix, a 10mm thick section of brain was dissected beginning ∼1 mm anterior to the cortical injury and extending caudally. (B) The 10 mm section was further dissected with a horizontal slice aligned with the bottom of the corpus callosum as shown, and a vertical slice at the midline to separate the left (contralesional) and right (injured or sham-injured) sides. The dorsal brain samples from each side were used for analysis of brain water content.

### MRI analysis

Regions of interest (ROIs) were drawn over the ipsilesional and contralesional cortex for each subject using MIPAV software. ROIs were drawn on three sequential diffusion weighted images (b = 50 s/mm^2^) encompassing the center of the injury (centered at bregma). First a vertical line was drawn at the midline. Then, from the point where the midline intersects the base of the brain, a second line was drawn at a 45-degree angle. The ROI was completed by connecting the points where these lines crossed the cortex, following the dorsal surface of the brain and the corpus callosum. After the ROI was drawn for the ipsilesional cortex, it was flipped 180 degrees across the vertical midline and manually adjusted to fit the contralesional cortex. The same ROIs, generated by a single experimenter, were used for analysis of both T_2_ and DWI data.

Because the ROIs encompassing the lesion contain a mix of lesion pixels, normal pixels, and in some cases bleeding that could influence the quantitative analysis, we used a thresholding approach to focus analysis on the more injured areas and exclude any potential confound from bleeding (**Supplementary Figure 1**). Using the T_2_ maps (where higher values correspond to pathology, middle values to normal cortical tissue, and lower values to bleeding), we generated histograms of all pixel intensities within the injured and contralesional ROIs. For each animal, we generated a gaussian curve to fit the normal distribution of pixels in the contralesional ROI and used to this to set a threshold (μ – σ). Image masks were generated of the ROI pixels above the threshold, and we performed quantitative analyses on the T_2_ and ADC values from the masked regions, averaged across the three image slices. Finally, we calculated the change in T_2_ or ADC observed in the injured ROI compared with the contralesional side: [(Injured side/Contralesional side)*100 – 100].

### Statistical analysis

We used two-way ANOVA to compare the hemispheric change in tissue water, T_2_, and ADC measures across group (TBI/sham) and sex (male/female), with follow-up by the Wilcoxon rank sum test to compare groups within each sex. Correlations between MRI parameters and brain water content were assessed using the Spearman test. All tests were two-sided and performed with R software^61^. Statistical significance was defined as *p* < 0.05. Due to the exploratory nature of the study and the relatively small group sizes, we did not control for multiple comparisons

### Subject exclusions and MRI quality screening

Fifteen male and 15 female rats received CCI; eight male and eight female rats received sham surgery. One female TBI rat died during imaging and was excluded from the study. One female sham rat was excluded from the study due to surgical error. Tissue water measurements were excluded from 1 female TBI and 1 female sham rat due to data collection error. Quality assessments of MRI data were performed by a blinded screener. For DWI, motion artifacts or data corruption led to exclusion of 2 male TBI, 2 male sham, 2 female TBI, and 2 female sham rats. For the T_2_ data, image artifacts or data corruption led to exclusion of 2 male TBI, 1 female TBI, and 1 female sham.

## AUTHOR CONTRIBUTIONS

Heather M Minchew: Conceptualization, Methodology, Investigation, Writing-Original draft;

Sadie L Ferren: Investigation, Data Curation, Visualization;

Sarah K Christian: Investigation, Resources;

Jinxiang Hu: Formal analysis, Writing-Reviewing and Editing;

Paul Keselman: Methodology, Software;

William M Brooks: Writing-Reviewing and Editing;

Brian T Andrews: Writing-Reviewing and Editing;

Janna L Harris: Conceptualization, Methodology, Investigation, Project administration, Funding acquisition, Writing-Reviewing and Editing.

## SUPPLEMENTARY FILES

**Supplementary Figure 1.**
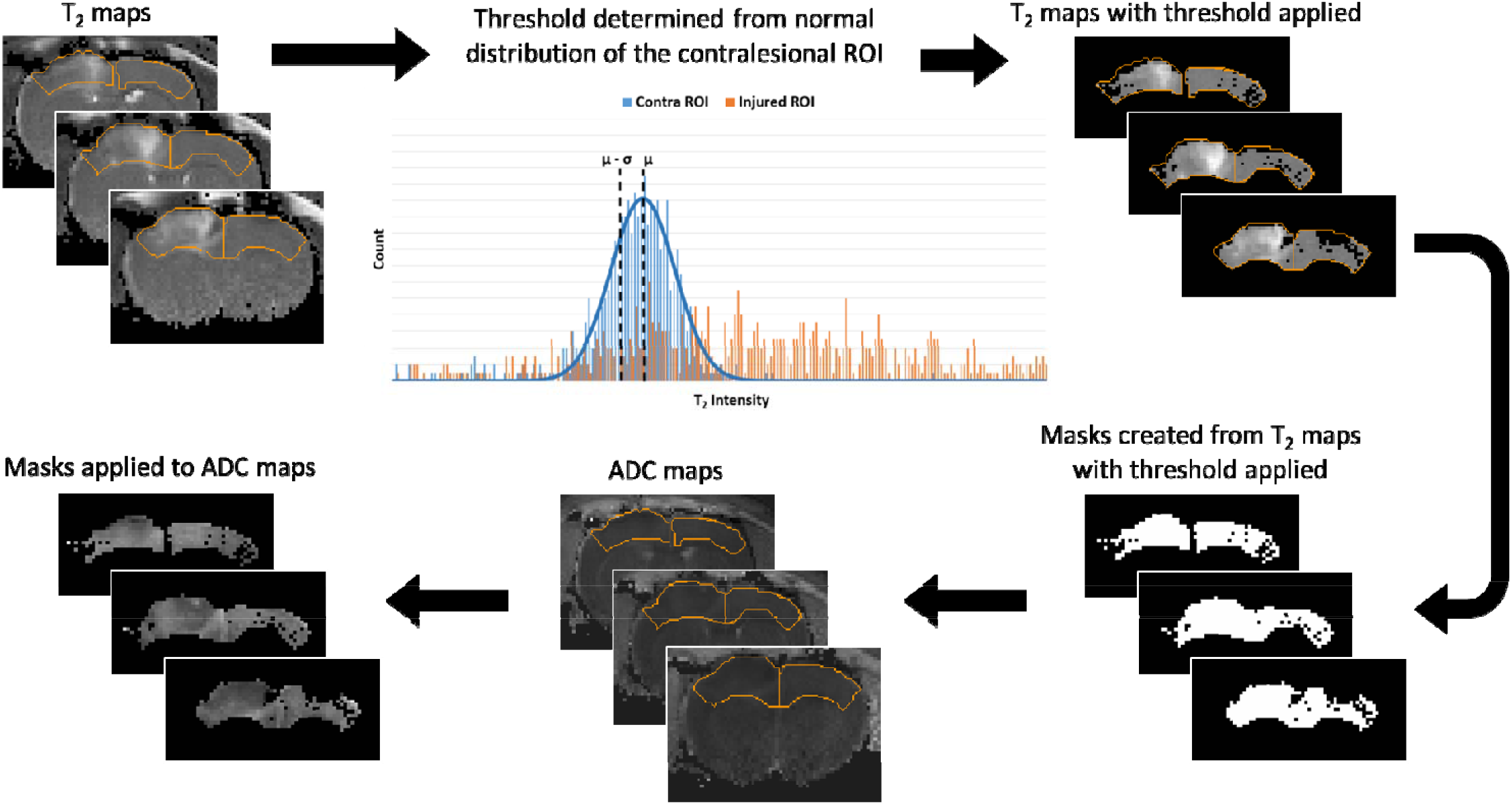
Thresholding protocol for analysis of T_2_ and ADC data.

**Supplementary Table 1.**
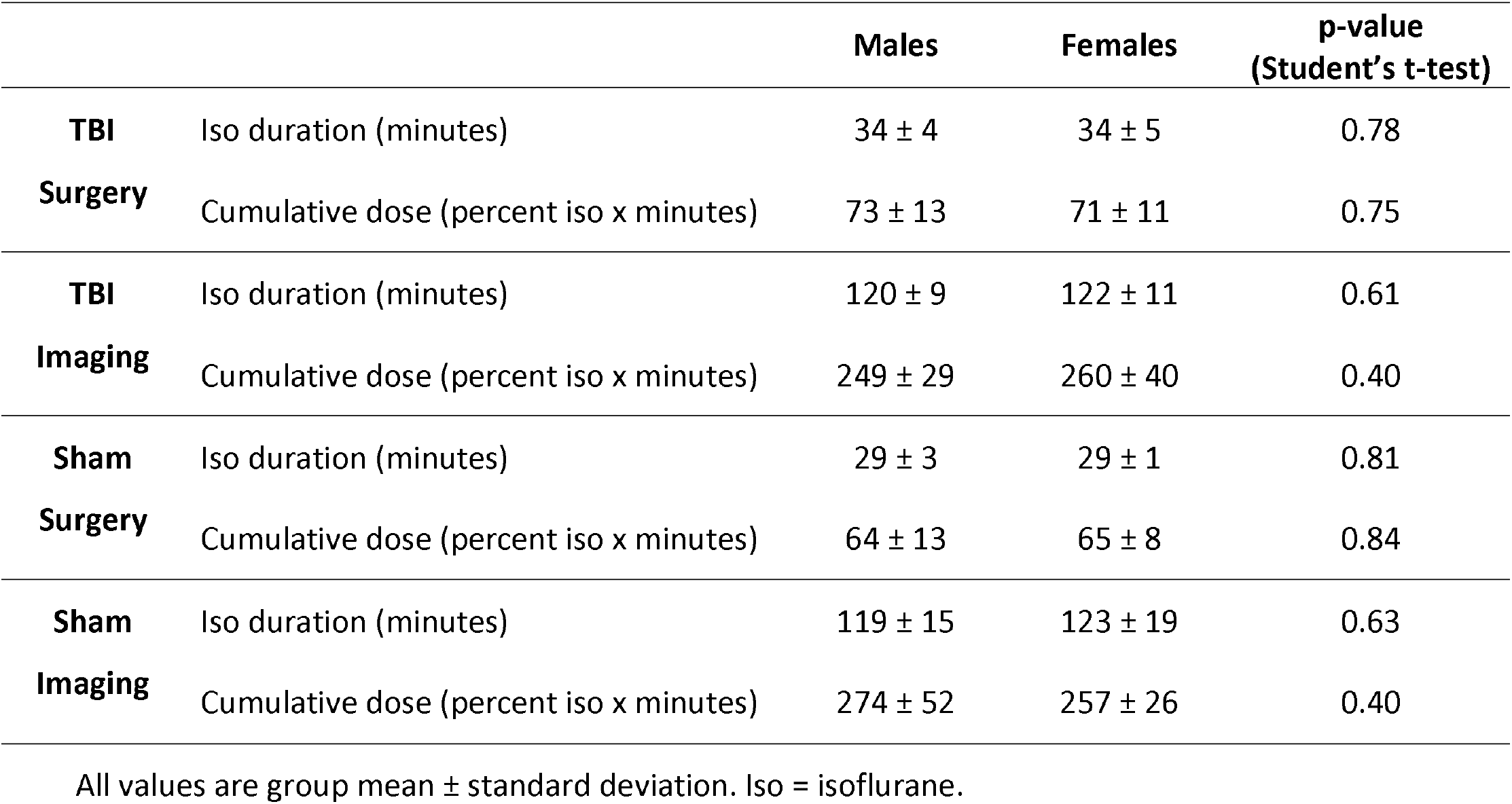
Isoflurane duration and estimated dosing.

